# Treadmilling by FtsZ filaments drives peptidoglycan synthesis and bacterial cell division

**DOI:** 10.1101/077560

**Authors:** Alexandre W. Bisson Filho, Yen-Pang Hsu, Georgia R. Squyres, Erkin Kuru, Fabai Wu, Calum Jukes, Cees Dekker, Seamus Holden, Michael S. VanNieuwenhze, Yves V. Brun, Ethan C. Garner

## Abstract

How bacteria produce a septum to divide in two is not well understood. This process is mediated by periplasmic cell-wall producing enzymes that are positioned by filaments of the cytoplasmic membrane-associated actin FtsA and the tubulin FtsZ (FtsAZ). To understand how these components act in concert to divide cells, we visualized their movements relative to the dynamics of cell wall synthesis during cytokinesis. We find that the division septum is built at discrete sites that move around the division plane. Furthermore, FtsAZ filaments treadmill in circumferential paths around the division ring, pulling along the associated cell-wall-synthesizing enzymes. We show that the rate of FtsZ treadmilling controls both the rate of cell wall synthesis and cell division. The coupling of both the position and activity of the cell wall synthases to FtsAZ treadmilling guides the progressive insertion of new cell wall, synthesizing increasingly small concentric rings to divide the cell.

**One-sentence summary:** Bacterial cytokinesis is controlled by circumferential treadmilling of FtsAZ filaments that drives the insertion of new cell wall.

## Main Text

Cells from all domains of life must divide in order to proliferate. In bacteria, cell division involves the inward synthesis of the cell wall peptidoglycan (PG), creating a septum that cleaves the cell in two. Septation is directed by proteins that are highly conserved among bacteria. The location and activity of the septal PG synthesis enzymes are regulated by FtsZ, a tubulin homolog, which associates with the cytoplasmic side of the membrane via an actin-like protein FtsA. FtsZ can form filaments (*1*) and membrane-associated copolymers with FtsA (FtsAZ) (*2*). Together, the two proteins form a dynamic structure, called the Z ring, that encircles the cell at the future division site (*3*), recruiting PG synthases and other proteins involved in cytokinesis (*4*, *5*). Once the division machinery is mature, the cytosolic Z ring constricts, while the associated enzymes on the other side of the membrane build a PG-containing septum that partitions the cell in two.

Whereas a detailed catalog of cell division proteins and their interdependency for septal localization is known, we do not have a clear picture of how these components interact with each other in space and time to carry out cytokinesis. One principal obstacle to understanding cytokinesis has been the inability to observe the dynamics of individual components relative to each other, and relative to the septal PG structure they build. For example: the organization and dynamics of FtsZ filaments within the Z ring remain ill defined, it is not known how FtsAZ filaments control the activity of PG synthases, and the pattern of septal PG synthesis during division has never been directly observed. To gain insight into how these components work together to divide bacteria, we visualized the dynamics of septal PG synthesis in relation to the movements of FtsAZ filaments and the septal PG synthase Pbp2B in the Gram-positive *Bacillus subtilis*.

To assess the dynamics of septal PG synthesis, we sequentially pulse-labeled growing cells with different colors of fluorescent D-amino acids (FDAAs), probes that incorporate into PG (*6*) by the D,D-transpeptidation activity of PG synthases (*7*, *8*). 3D-structured illumination microscopy (3D-SIM) analysis showed that sequential three color pulse labeling resulted in bullseye patterns at the division plane, with the first FDAA on the outside, the second FDAA in the middle, and the last at the center (Fig. 1A). We conclude that the septum is progressively synthesized inward from the cell surface, in agreement with early electron microscopy studies (*9*). In contrast to long pulses, short sequential pulses of two FDAA colors resulted in discrete spots or arcs distributed around the septum, with the 2 colors offset from each other. Cells labeled with sequential FDAA pulses had decreased colocalization of FDAA incorporation compared to cells labeled with two colors simultaneously (Fig. 1B-C, **S1A**). Therefore, PG synthesis occurs at discrete sites that move around the division plane.

**Figure 1:**
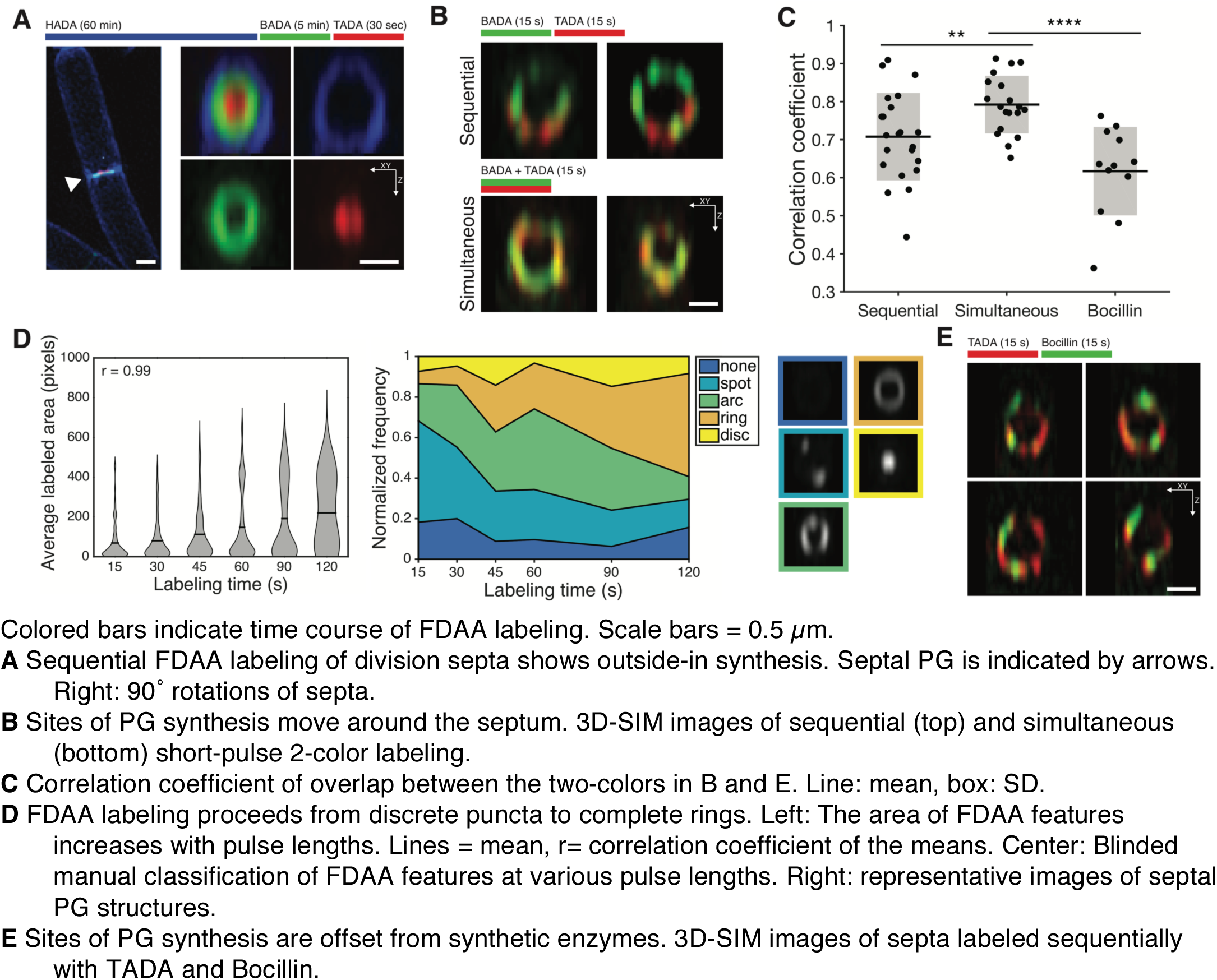
Septal PG synthesis occurs at discrete, mobile sites.

We then tested how discrete sites of PG synthesis evolve into a complete division septum by labeling cells with FDAAs using increasing pulse durations. The area of the labeled spots and arcs increased with the pulse duration, as did the total amount of labeling at the septum (Fig. 1D, left; **Fig. S2A-C**). With short pulses (15 sec), cells had discrete spots or arcs of FDAA label (**Fig. S1B**). As pulse duration increased, cells displayed an increased frequency of arcs, gradually transitioning into complete rings at longer pulse lengths (Fig. 1D, **right**). PG synthesis inhibitors greatly reduced FDAA incorporation, indicating that labeling was specific (**Fig. S2D**). To explore where the PG synthesis enzymes were located relative to newly incorporated PG, we followed a short FDAA pulse by a treatment with Bocillin, which both inactivates and fluorescently labels PBPs (Fig. 1E). The Bocillin signal only partially colocalized with the newly-synthesized PG (Fig. 1C). Bocillin has been proposed (*10*, *11*) to label active PBPs, so we infer that the PG synthases might move around the division plane.

To explore the possibility that PG is processively synthesized around the division plane, we examined the motions of the division-specific PG synthases and associated cytoskeletal polymers. Total internal reflection microscopy (TIRFM) of a functional mNeonGreen-FtsZ fusion expressed from the native locus (**Fig. S3A-C**) frequently revealed directional movements of FtsZ signal within newly assembled Z rings (Fig. 2A, **Movie SM1**). Furthermore, in almost every cell, we observed small filaments outside the Z ring moving directionally along the short axis at the same rate as the motion observed within Z rings. Overexpression of a second chromosomally integrated *ftsAZ* operon (FtsA, mNeonGreen-FtsZ) resulted in many more directionally moving filaments outside the Z ring (Fig. 2E, **S3D, Movie SM1**). To resolve FtsZ motion in mature Z rings, where the dense signal made directional m ovement difficult to observe, we vertically immobilized bacteria in agarose microholes, orienting the division plane parallel to the objective (**Fig. S4**). Widefield imaging revealed multiple FtsZ filaments moving in both directions around the circumference of the constriction site (Fig. 2C, **Movie SM2**) over a wide range of ring diameters (400-800 nm), confirming that FtsZ filaments move directionally around dense, actively constricting rings. Movement of FtsZ filaments around the Z ring explains the heterogeneous, dynamic structures obtained via super resolution microscopy (*12*–*16*), as well as the fast turnover of its internal subunits (*17*). Motions identical to FtsZ were observed with a functional FtsA-mNeonGreen fusion (Fig. 2B, **Movie SM1**), and two-color imaging of FtsA and FtsZ indicated that they colocalize and move together (Fig. 2D). We conclude that filaments of FtsAZ move directionally around the division plane.

**Figure 2:**
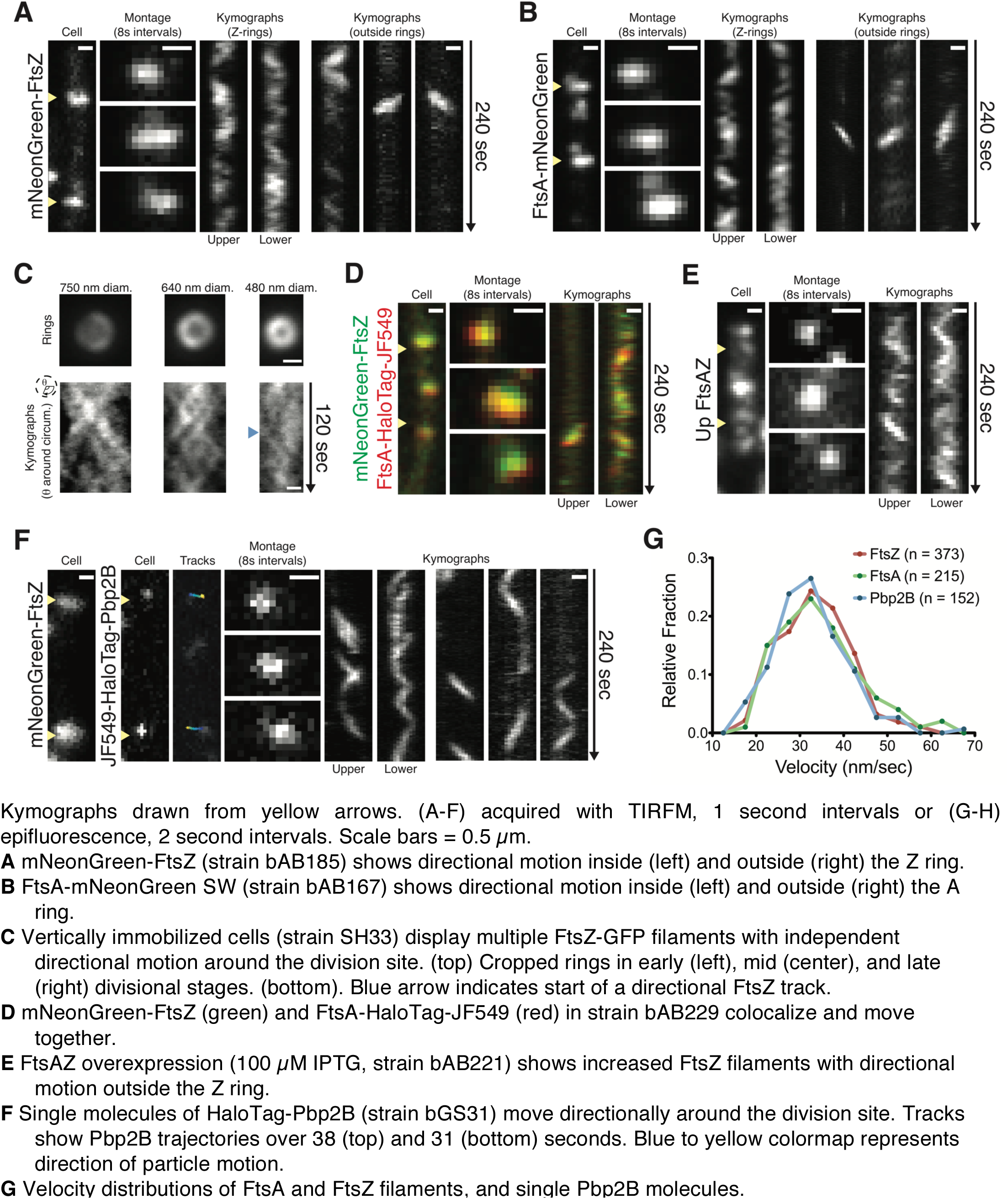
FtsAZ and Pbp2B move directionally around the division site.

This led us to ask if the division-associated transpeptidase Pbp2B moves with FtsAZ. At native expression levels, mNeonGreen-Pbp2B moved directionally along the Z ring axis (**Fig. S5A**); these motions could be observed more clearly at reduced expression levels (**Fig. S5B**). To observe the motions of single Pbp2B molecules, we used low concentrations of HaloLigand-JF549 (*18*) to label HaloTag-Pbp2B expressed from the native locus. TIRF imaging revealed two types of Pbp2B motion: 1) rapid diffusion within the membrane, and 2) directional, processive motions of Pbp2B moving around the cell width, always localized to Z rings (Fig. 2F, **S5C-E, Movie SM3**). In some cases, multiple Pbp2B molecules were observed moving directionally within the same Z ring, sometimes in opposite directions. FtsZ, FtsA, and Pbp2B all move at a similar velocity (Fig. 2G, **S5D**), suggesting their motions are associated. Combined with the apparently mobile nature of septal PG synthesis shown above, the directional movements of FtsAZ/Pbp2B around the division plane suggest that FtsAZ circumferential motions could be coupled to the insertion of new PG.

Next, we investigated the mechanism driving FtsAZ/Pbp2B motion. The coupled, circumferential motions of FtsAZ/Pbp2B appear similar to MreB/Pbp2A, the filament/enzyme system required for elongation of rod-shaped bacteria wherein the enzymatic activity of Pbp2A is required for MreB motion (*19*–*21*). Accordingly, we tested whether inhibition of Pbp2B would halt FtsAZ motion. However, FtsAZ velocity was unaffected by high concentrations of multiple PG synthesis inhibitors (Fig. 3A, 3H, **Movie SM4**) or by depleting cells of Pbp2B (Fig. 3B, 3H, **Movie SM5**).

**Fig 3:**
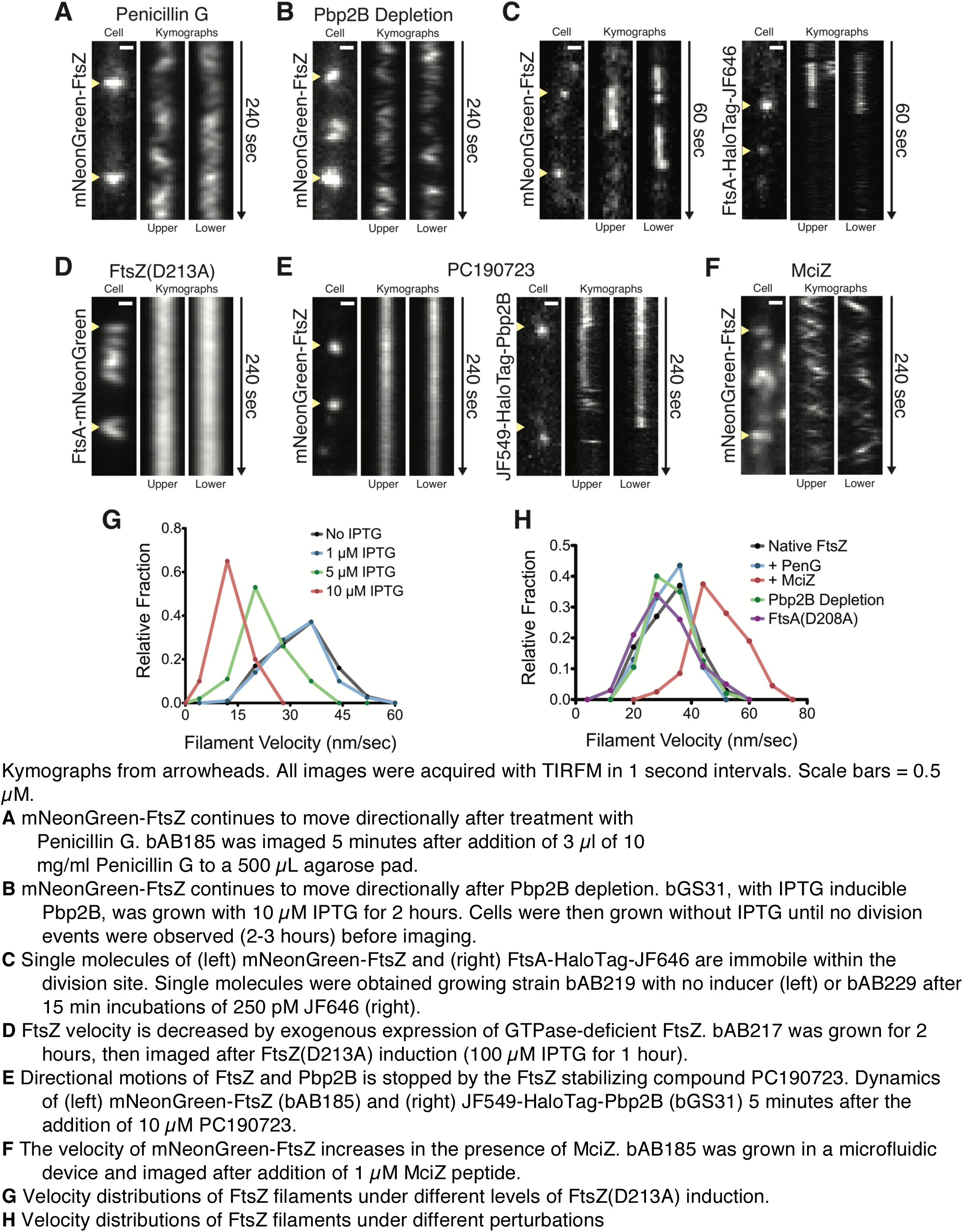
Directional motion of FtsAZ is driven by treadmilling, independent of cell wall synthesis, and required for Pbp2B motion.

We then tested if directional FtsAZ motion is due to filament treadmilling, as observed for FtsAZ assembled upon lipid bilayers *in vitro* (*22*). Consistent with treadmilling, sparse labeling of FtsZ or FtsA in cells demonstrated that single molecules of both proteins are immobile within moving filaments (Fig. 3C, **Movie SM6**). As treadmilling requires nucleotide hydrolysis (*23*), we measured the rate of FtsA filament motion as we modulated the nucleotide hydrolysis of FtsA and FtsZ. In agreement with *in vitro* studies (*22*), FtsA deficient in ATP hydrolysis displayed normal dynamics (Fig. 3H, **Movie SM7A**). In contrast, exogenous expression of FtsZ(D213A), a mutant with a greatly reduced GTPase rate (*24*), gradually reduced FtsAZ velocity, stopping motion at high inductions (Fig. 3D, 3G, **Movie SM7B**). Likewise, addition of PC190723, an inhibitor of FtsZ GTP hydrolysis (*25*) halted FtsZ movement (Fig. 3E, **Movie SM8A**). Conversely, expression of MciZ, a protein that increases the rate of FtsZ filament turnover (*26*), increased FtsZ velocity (Fig. 3F, 3H, **Movie S7C**). We next tested how FtsZ treadmilling dynamics affect Pbp2B movement. Stopping FtsAZ motion with PC190723 caused Pbp2B molecules to become immobile while remaining colocalized with FtsZ (Fig. 3E, **Movie SM8B**). Thus, FtsZ treadmilling drives the motions of FtsAZ filaments and the septal PG synthesis enzymes.

To understand how the directional motions of FtsAZ/Pbp2B relate to septal PG synthesis, we labeled cells with FDAAs as we altered FtsZ dynamics. Overexpression of FtsZ(D213A) created long, slowly-growing FtsA-mNeonGreen SW spirals (Fig. 4A, **Movie SM9**), and sequential FDAA labeling showed sequential incorporation along the entire spiral length (Fig. 4A). Likewise, long PC190723 treatments resulted in fragmented patches of both FtsZ and FDAA incorporation (**Fig. S6A**), indicating Pbp2B activity is constrained by FtsAZ location. However, these strong inhibitions of FtsZ dynamics required much longer pulse times to achieve FDAA labeling, suggesting that FtsAZ motion might be rate limiting for PG synthesis. To test this hypothesis, we altered FtsAZ velocity as we pulse-labeled cells with FDAAs, finding that both the amount and spatial distribution of PG synthesis within the ring correlate with FtsAZ velocity: conditions that slowed dynamics (D213A expression or PC190723) decreased both the total amount (Fig. 4B) and area (**Fig. S6B-C**) of FDAA labeling. Conversely, increasing FtsAZ velocity (MciZ) increased both the amount and area of labeling. Thus, septal PG synthesis depends on, and is limited by, the treadmilling of FtsZ filaments, rather than their presence.

**Fig 4:**
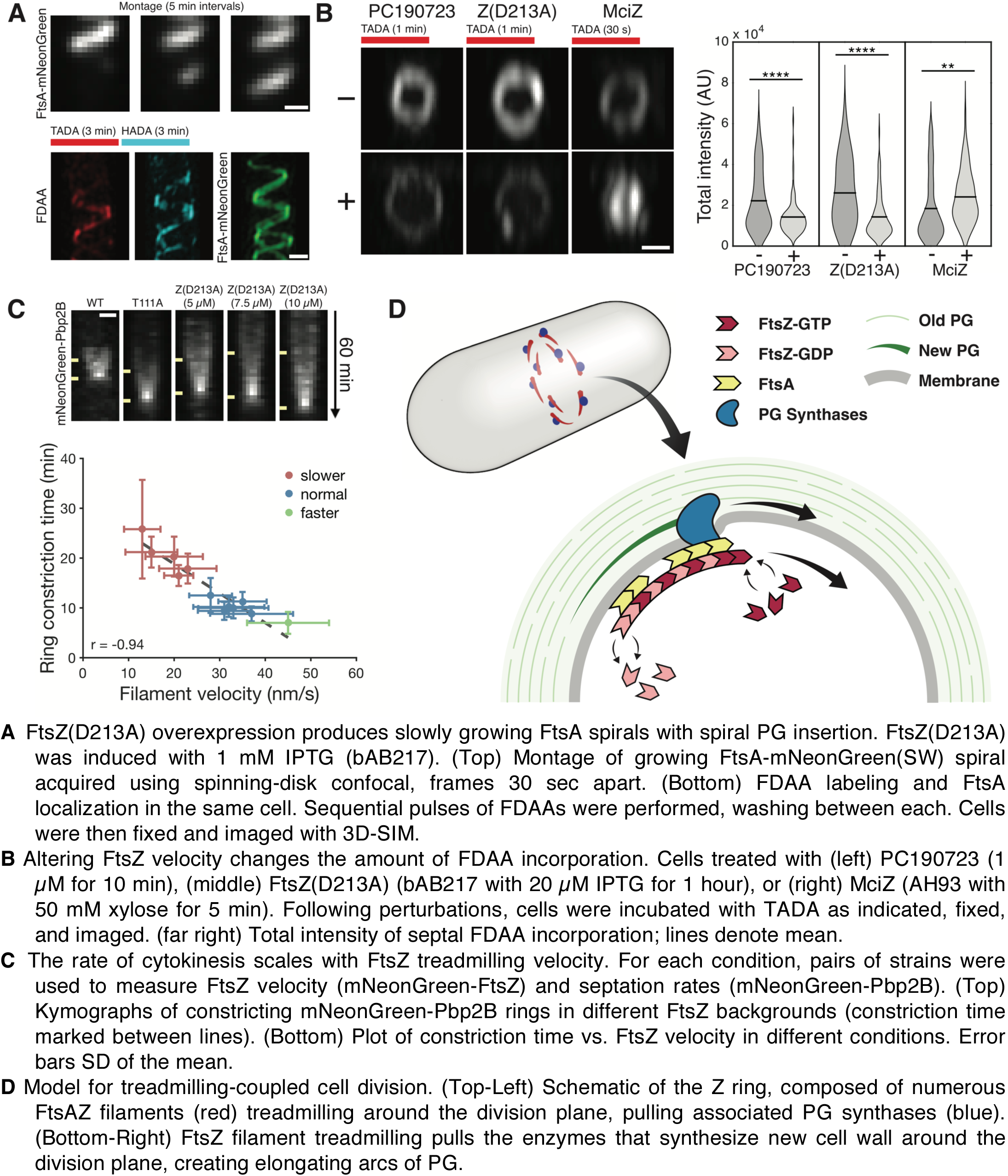
Cytokinesis is controlled by directional motion of FtsAZ filaments.

As these results indicate that FtsAZ dynamics control the rate of septal PG synthesis, we asked whether the rate of cytokinesis correlated with the rate of FtsAZ treadmilling. We modulated FtsAZ treadmilling velocity by 1) introducing different mutations reported to affect GTP hydrolysis into FtsZ at its native locus, 2) titrating expression of exogenous FtsZ(D213A), 3) MciZ expression, and 4) other perturbations (Fig. 4C, **S7A, Table S1, Movie SM10**). This analysis revealed that the rate of cytokinesis shows a linear dependence on FtsAZ velocity: division is slower as velocity decreases, and faster when velocity is increased. Even under the strongest perturbations of FtsZ hydrolysis, the decreased rates of cytokinesis are not due to decreased growth rates (**Fig. S7B**) as previously suggested (*27*). Combined, these experiments demonstrate that FtsAZ treadmilling dynamics are both linked to, and limiting for, septal PG synthesis and cell constriction in *B. Subtlilis*.

How bacteria coordinate cell wall synthesis to divide in two has remained unknown for 25 years after the discovery of Z ring. In sum, our results indicate that cell division occurs by the action of discrete enzyme-filament complexes, which, driven by FtsZ filament treadmilling, move around the division plane, building new PG during their transit (Fig. 4D). The treadmilling of FtsZ creates long range order out of the local activity of the PG synthases, linking circumferential enzyme motion to their insertion of new cell wall. This coupling may yield uniform insertion of new material around the division plane, causing the septum to be built inward in progressively smaller concentric rings (Fig. 1A), a pattern observed in *B. subtilis* septa (*28*).

The coupling between FtsZ treadmilling and PG synthesis can unifiy previously conflicting models of cell division. FtsZ filaments have been proposed to act as force generators to bend membranes (*2*, *29*), and as inert scaffolds for PG synthesis (*12*, *30*). If FtsZ filaments deform membranes, coupling their movement to PG synthesis would allow each local deformation to be reinforced via synthesis of PG into the gap between the membrane and old cell wall (*31*). These multiple sites of local deformation and their coupled reinforcing synthesis would then move around the division site, iteratively building the invaginating septum.

Interestingly, our findings reveal that the two essential and most conserved modes of bacterial growth – bacterial elongation and the building of the division septum – have evolved inverted mechanisms to control the nanoscale distribution of their PG synthesis. During elongation, PG synthesis enzymes pull stable MreB filaments (*19*–*21*). In contrast, during division, treadmilling FtsZ filaments pull the PG synthesis enzymes in addition to controling their activity, a novel role for cytomotive filaments.

## Author contributions

This work represents the merged research directions of 3 groups (EG, YB/MV, SH/CD). FDAA labeling and 3D SIM were performed by YH and EK. Strains were constructed by AB, GS, and CJ. Confocal, TIRF, and single molecule imaging were done by AB and GS. Imaging in vertical microholes was performed by SH and CJ. Programming for analysis was done by GS and SH. Microholes were designed by SH, CD, and FW. Microfabrication was conducted by FW. All authors designed experiments and wrote the paper.

## Acknowledgments

We thank F. Gueiros-Filho, R. Losick, M. Erb, and P. Levin for strains; L. Lavis for his gift of JF dyes; B. Murphy and E. Pasciak for help with FDAA synthesis; R. Losick, B. LaSarre, and D. Kearns for critical comments on the manuscript. This work was supported by NIH grants GM113172 to MSV and YVB; and GM51986 to YVB; a Searle Scholar Fellowship and NIH grant DP2AI117923-01 to ECG; a Newcastle University Research Fellowship and Royal Society Research Grant RG150475 to SH; a Science Without Borders Research Fellowship to AWB-F; an ERC Advanced Grant SynDiv 669598 to CD, and an NSF GRFP (DGE1144152) to GRS. SIM was performed in the Indiana University Light Microscopy Imaging Center supported by S10RR028697- 01.

## References

1. Y. Chen, H. P. Erickson, Rapid in vitro assembly dynamics and subunit turnover of FtsZ demonstrated by fluorescence resonance energy transfer. Journal of Biological Chemistry. 280, 22549–22554 (2005).

2. P. Szwedziak, Q. Wang, T. A. M. Bharat, M. Tsim, J. Lowe, Architecture of the ring formed by the tubulin homologue FtsZ in bacterial cell division. Elife. 3, 642 (2014).

3. X. Ma, D. W. Ehrhardt, W. Margolin, Colocalization of cell division proteins FtsZ and FtsA to cytoskeletal structures in living Escherichia coli cells by using green fluorescent protein. Proc Natl Acad Sci USA. 93, 12998–13003 (1996).

4. P. Gamba, J. W. Veening, N. J. Saunders, L. W. Hamoen, R. A. Daniel, Two-step assembly dynamics of the Bacillus subtilis divisome. J Bacteriol. 191, 4186–4194 (2009).

5. B. Söderström et al., Coordinated disassembly of the divisome complex in Escherichia coli. Mol Microbiol. 101, 425–438 (2016).

6. E. Kuru et al., In Situ probing of newly synthesized peptidoglycan in live bacteria with fluorescent D-amino acids. Angew. Chem. Int. Ed. Engl. 51, 12519–12523 (2012).

7. T. J. Lupoli et al., Transpeptidase-mediated incorporation of D-amino acids into bacterial peptidoglycan. J Am Chem Soc. 133, 10748–10751 (2011).

8. Y. Qiao et al., Detection of Lipid-Linked Peptidoglycan Precursors by Exploiting an Unexpected Transpeptidase Reaction. J Am Chem Soc, 141010140136004 (2014).

9. J. Hillier, J. F. Hoffman, On the ultrastructure of the plasma membrane as determined by the electron microscope. Journal of cellular physiology. 42, 203–247 (1953).

10. C. Eberhardt, L. Kuerschner, D. S. Weiss, Probing the catalytic activity of a cell division-specific transpeptidase in vivo with beta-lactams. J Bacteriol. 185, 3726–3734 (2003).

11. O. Kocaoglu et al., Selective Penicillin-Binding Protein Imaging Probes Reveal Substructure in Bacterial Cell Division. 7, 8 (2012).

12. C. Coltharp, J. Buss, T. M. Plumer, J. Xiao, Defining the rate-limiting processes of bacterial cytokinesis. Proc Natl Acad Sci USA. 113, E1044–53 (2016).

13. S. J. Holden et al., High throughput 3D super-resolution microscopy reveals Caulobacter crescentus in vivo Z-ring organization. Proc Natl Acad Sci USA. 111, 4566–4571 (2014).

14. M. P. Strauss et al., 3D-SIM Super Resolution Microscopy Reveals a Bead-Like Arrangement for FtsZ and the Division Machinery: Implications for Triggering Cytokinesis. PLoS Biol. 10 (2012), doi:10.1371/journal.pbio.1001389.

15. H.-C. T. Tsui et al., Pbp2x localizes separately from Pbp2b and other peptidoglycan synthesis proteins during later stages of cell division of Streptococcus pneumoniae D39. Mol Microbiol. 94, 21–40 (2014).

16. M. Jacq et al., Remodeling of the Z-Ring Nanostructure during the Streptococcus pneumoniae Cell Cycle Revealed by Photoactivated Localization Microscopy. mBio. 6, e01108–15 (2015).

17. D. E. Anderson, F. J. Gueiros-Filho, H. P. Erickson, Assembly dynamics of FtsZ rings in Bacillus subtilis and Escherichia coli and effects of FtsZ-regulating proteins. J Bacteriol. 186, 5775–5781 (2004).

18. J. B. Grimm et al., A general method to improve fluorophores for live-cell and single-molecule microscopy. Nat Meth. 12, 244–250 (2015).

19. E. C. Garner et al., Coupled, circumferential motions of the cell wall synthesis machinery and MreB filaments in B. subtilis. Science. 333, 222–225 (2011).

20. J. Domínguez-Escobar et al., Processive Movement of MreB-Associated Cell Wall Biosynthetic Complexes in Bacteria. Science. 333, 225–228 (2011).

21. S. van Teeffelen et al., The bacterial actin MreB rotates, and rotation depends on cell-wall assembly. Proc Natl Acad Sci USA. 108, 15822–15827 (2011).

22. M. Loose, T. J. Mitchison, The bacterial cell division proteins FtsA and FtsZ self-organize into dynamic cytoskeletal patterns. Nat Cell Biol. 16, 38–46 (2014).

23. J. A. Theriot, The polymerization motor. Traffic. 1, 19–28 (2000).

24. S. D. Redick, J. Stricker, G. Briscoe, H. P. Erickson, Mutants of FtsZ targeting the protofilament interface: effects on cell division and GTPase activity. J Bacteriol. 187, 2727–2736 (2005).

25. D. J. Haydon et al., An inhibitor of FtsZ with potent and selective anti-staphylococcal activity. Science. 321, 1673–1675 (2008).

26. A. W. Bisson-Filho et al., FtsZ filament capping by MciZ, a developmental regulator of bacterial division. Proc Natl Acad Sci USA. 112, E2130–8 (2015).

27. C. Coltharp, J. Buss, T. M. Plumer, J. Xiao, Defining the rate-limiting processes of bacterial cytokinesis. Proc Natl Acad Sci USA. 113, E1044–E1053 (2016).

28. E. J. Hayhurst, L. Kailas, J. K. Hobbs, S. J. Foster, Cell wall peptidoglycan architecture in Bacillus subtilis. Proc Natl Acad Sci USA. 105, 14603–14608 (2008).

29. M. Osawa, H. P. Erickson, Liposome division by a simple bacterial division machinery. Proc Natl Acad Sci USA. 110, 11000–11004 (2013).

30. A. J. F. Egan, W. Vollmer, The stoichiometric divisome: a hypothesis. Front Microbiol. 6, 0–6 (2015).

31. Z. Li, M. J. Trimble, Y. V. Brun, G. J. Jensen, The structure of FtsZ filaments in vivo suggests a force-generating role in cell division. EMBO J. 26, 4694–4708 (2007).

